# Decoding commensal-host communication through genetic engineering of *Staphylococcus epidermidis*

**DOI:** 10.1101/664656

**Authors:** Y. Erin Chen, Nicolas Bouladoux, Charlotte Hurabielle, Aiden M. Mattke, Yasmine Belkaid, Michael A. Fischbach

## Abstract

Commensal skin bacteria elicit potent, antigen-specific immune responses in the skin without barrier breach or visible inflammation. While microbial modulation of immune homeostasis has profound consequences for epithelial health and inflammatory skin diseases, the mechanisms of microbe-immune crosstalk in the skin are largely unknown. A key barrier to mechanistic work has been genetic intractability of one of the most prevalent skin colonists, *Staphylococcus epidermidis* (*S. epidermidis*). Here, we develop a novel method to create a library of mutants with defined cell envelope alterations in primary human *S. epidermidis* isolates. By colonizing mice with these mutants, we uncover bacterial molecules involved in the induction of defined immune signatures. Notably, we show that under conditions of physiologic colonization, *S. epidermidis* cell envelope glycolipids are sensed by C-type lectin receptors, likely in non-myeloid cells, in conjunction with Toll-like receptors. This combinatorial signaling determines the quality of T cell responses and results in the potential for greater specificity toward commensal microbiota than previously appreciated. Additionally, the microbial molecules required for the colonization-induced immune response are dispensable for T cells responses in a model of *S. epidermidis* infection, but differentially modulate innate inflammatory responses. Thus, the same microbe uses distinct sets of molecules to signal to the immune system commensal versus pathogenic behavior, and differential sensing of these microbial signals depends on host context.

## MAIN TEXT

Commensal skin bacteria elicit potent, taxon-specific immune responses in the skin under conditions of normal colonization, without breach of the skin barrier (Naik et al., 2012; Ridaura et al., 2018). These microbiota-derived stimuli play a critical role in reactivity to inflammatory triggers (Ridaura et al., 2018; Scharschmidt et al., 2015, 2017), pathogen resistance (Naik et al., 2015), and wound healing (Harrison et al., 2019; Linehan et al., 2018; Williams et al., 2019). Despite their fundamental importance to skin biology, the mechanisms by which commensals and immune cells interact are poorly understood. The unique features of commensal sensing include: (1) presumptive sampling of whole microbial cells (or microbial products) through an intact epithelial barrier, (2) a low biomass of highly diverse microbial signals relative to infection, and (3) the lack of inflammatory signals that would result from epithelial injury. These features make sensing of commensal organisms and orchestrating a balanced, homeostatic immune response a unique process, distinct from the sensing of an invading pathogen. Defining the mechanisms of microbe-immune cell communication would allow us to understand the molecular underpinnings of immunity at the skin barrier and identify new targets for modulating immune responses in the context of skin disease.

Most efforts to study commensal-host interactions have focused on the host genes and cell populations involved, rather than the microbial molecules responsible for driving host responses. In the case of *S. epidermidis*, colonization provokes a non-inflammatory CD4^+^ and CD8^+^ T cell response with wound healing and pathogen resistance properties in both mice and non-human primates (Linehan et al., 2018; Naik et al., 2012, 2015). In particular, the CD8^+^ T cell response is characterized by the co-expression type 17 and type 2 programs, allowing these cells to promote both antimicrobial defense and wound repair (Harrison et al., 2019; Linehan et al., 2018; Naik et al., 2012, 2015). *S. epidermidis*-induced CD8^+^ T cells are specific for *S. epidermidis* N-formylated peptide antigens that are presented by nonclassical MHC I molecule H2-M3, a process that depends upon dermal dendritic cells (Linehan et al., 2018; Naik et al., 2015). Although the class of cognate peptide antigens is now known for CD8^+^ T cells, the *S. epidermidis* molecules that direct epithelial and antigen-presenting cell signaling in the initial stages of encounter, ultimately directing the downstream CD8^+^ and CD4^+^ T cell response, are undefined. More generally, the mechanisms and molecular partners involved in host-microbiota interaction remain poorly understood.

A major barrier to mechanistic understanding of this *S. epidermidis*-immune interaction has been the genetic intractability of *S. epidermidis*, a limitation common to most members of the human microbiota. *S. epidermidis* colonizes the skin of every human (Grice et al., 2009), and in the context of barrier breach (for example, device implantation) or immunocompromise, it is a frequent pathogen (Otto, 2009). However, only a few reports exist of methods for targeted mutations in *S. epidermidis* (Brophy et al., 2018; Monk et al., 2012; Winstel et al., 2016). Previously described genetic methods have advanced the field, but they are time-consuming and effective only for a few specific strains of *S. epidermidis* (Monk et al., 2012; Winstel et al., 2016) or able to make insertions only at specific genomic sites (Brophy et al., 2018). Here, we introduce a new method to genetically engineer *S. epidermidis* that is efficient and works in the vast majority of primary human *S. epidermidis* isolates. Driven by the observation that specific host pattern recognition receptor (PRR) mutants altered or abolished components of the adaptive immune response to *S. epidermidis*, we used this method to remove or alter components of the *S. epidermidis* cell wall, which is a rich source of candidate PRR ligands.

Under conditions of physiologic colonization, we found that *S. epidermidis* cell envelope mutants elicit marked changes in the composition and function of the T cell population in the skin. Modifications to the lipid versus carbohydrate moieties of surface structures yield distinct and unexpected shifts in the T cell response due to sensing of microbial components by C-type lectin receptors and Toll-like receptors. Our data reveal two unexpected findings. First, *S. epidermidis* cell wall ligands are sensed combinatorially by host receptors, creating the potential for much greater specificity and control of immune output toward commensal strains than previously appreciated. Second, while mutants in the *S. epidermidis* envelope elicit marked differences in the T cell number and quality during colonization, they do not alter the intensity of the T cell response to *S. epidermidis* intradermally, while impacting defined innate cell responses. These data indicate that the molecular components of the *S. epidermidis* envelope are ‘commensalism factors’ that guide the homeostatic immune response during colonization but not the anti-infection immune response in the context of invasion and breach of the skin barrier. These findings establish a molecular mechanism for homeostatic immune induction by *S. epidermidis,* and they open the door to exogenous control of the host pathways governing immune modulation by skin commensals.

### A method for genetically modifying primary *S. epidermidis* isolates

Targeted genetic modification of *S. epidermidis* has been challenging for two reasons. First, *S. epidermidis* has multiple stringent restriction systems that differ substantially among strains (Lee et al., 2016; Monk et al., 2012, 2015). Among human commensal skin bacteria, *S. epidermidis* has a relatively large pan-genome with a high degree of genetic variation among individuals (20% variability compared to 5% for *Cutibacterium acnes*) (Oh et al., 2016). As a result, genetic methods developed for *Staphylococcus aureus* that rely on bypassing strain-specific restriction-modification systems (Monk et al., 2015) are difficult to apply to *S. epidermidis*. Second, like many other Gram-positive bacteria, *S. epidermidis* can only be electroporated inefficiently due to a thick cell wall (Moran et al., 2018). Published methods demonstrate a transformation efficiency in *S. epidermidis* strain RP62a of approximately 20 cfu/μg of plasmid DNA compared to 10^3^ cfu/μg for *S. aureus* (Monk et al., 2012) and 10^9^ cfu/μg for *E. coli* using a standard protocol (Inoue et al., 1990).

To bypass poor electroporation efficiency, a genetic approach was recently described that involves transduction with bacteriophage Φ187 from an engineered restriction-deficient S*. aureus* (PS187 Δ*hsdR* Δ*sauPSI*) to *S. epidermidis* or other coagulase-negative *Staphylococcus* species on the basis of their similarity in wall teichoic acid structure (Winstel et al., 2013, 2016). Although this method works well for a specific clade of *S. epidermidis* strains, it is time-consuming, requires prolonged phage propagation, and is useful only for certain strains of *S. epidermidis* with efficient phage adsorption capacity; for example, this method does not work for *S. epidermidis* ATCC 12228. Given the high variability of phage specificity across *S. epidermidis* strains, the application of this phage-based method across many primary human isolates of *S. epidermidis* is limited, thus restricting our ability to study strain-strain and strain-host interactions.

Here, we introduce a novel electroporation and heat shock-based method that takes advantage of the methylase-deficient (Δ*dcm*) *E. coli* strain DC10B (Monk et al., 2012) and includes additional measures that increase electroporation efficiency and decrease restriction-modification efficiency. Critical elements of our method include cultivating *S. epidermidis* in hyperosmolar sorbitol, washing thoroughly in high volume 10% glycerol, a heat shock before or after electroporation, and a prolonged recovery period prior to plating (**Figure 1A**). Our approach does not restrict the efficiency of cloning to phage type and does not require specialized knowledge of strain-specific restriction systems. Using this method, we were able to construct mutations in most of the *S. epidermidis* strains we tested, include >10 primary human isolates from diverse phylogenetic groups (**Figure 1B-C**).

**Fig. 1.**
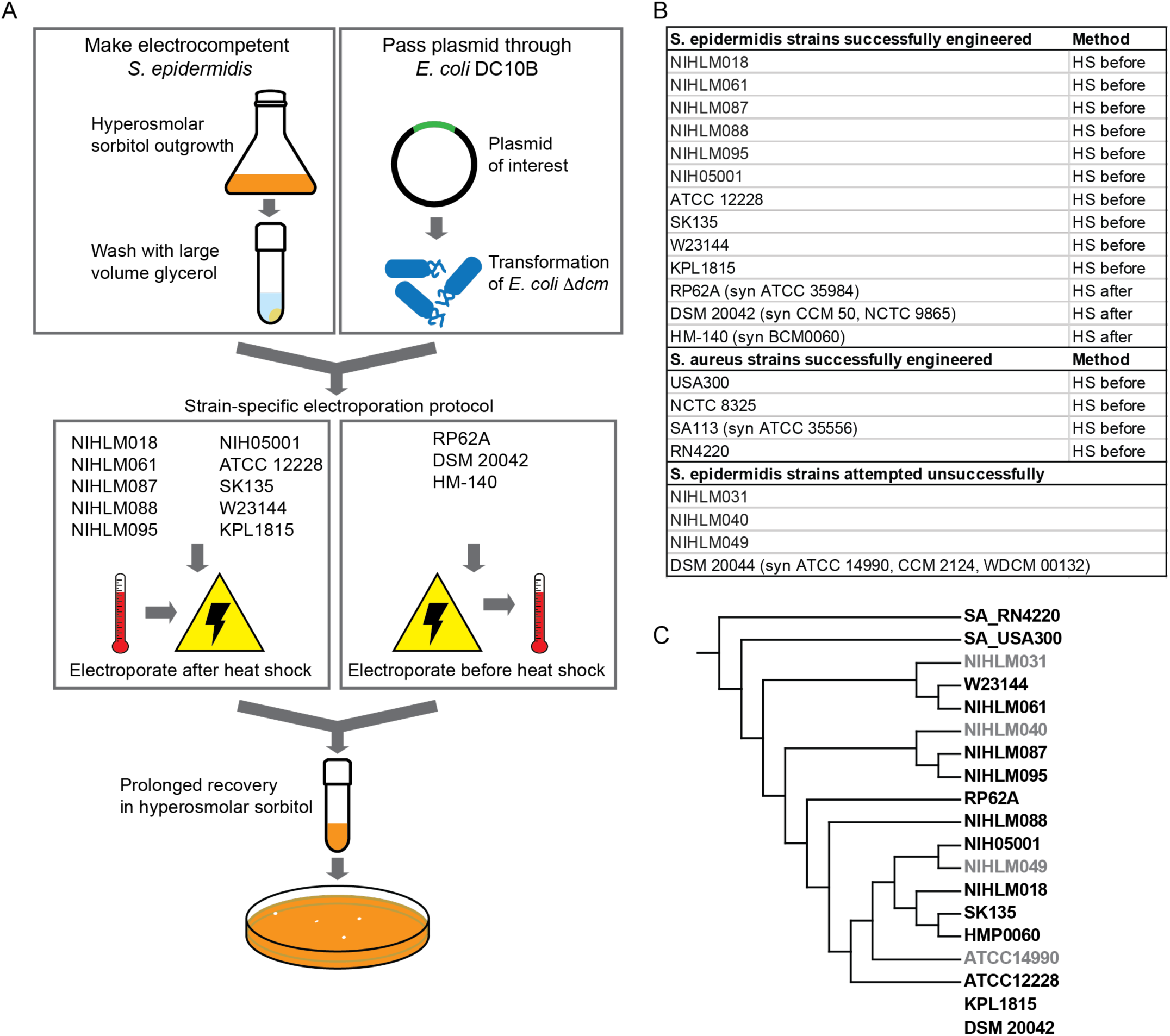
A new method for genetic modification of *S. epidermidis*. (A) Schematic of *S. epidermidis* genetic engineering method. To optimize electroporation efficiency, the *Staphyloccocus* strain is grown in media containing hyperosmolar sorbitol, harvested during late-log phase, and thoroughly washed with a large volume of 10% glycerol to eliminate salts. To bypass stringent restriction systems, the target plasmid is passed through *E. coli* Δ*dcm* and *Staphylococcus* is subjected to heat shock before or after electroporation. (B) Table of *S. epidermidis* and *S. aureus* strains that have been genetically modified in this work. The two methods differ only in the timing of the heat shock: prior to electroporation (“HS before”) or in the recovery medium after electroporation (“HS after”). Our method enables genetic modification of 13/17 *S. epidermidis* strains attempted. 4/4 *S. aureus* strains attempted were modified, including those that are restriction-competent. (C) Phylogenetic tree of *S. epidermidis* and *S. aureus* strains demonstrating that our method works across diverse *S. epidermidis* strains, including two strains that do not have genome sequences available in NCBI. Black = genetically manipulable. Grey = not genetically manipulable with our method.

### Specific pattern recognition receptors drive the adaptive immune response to *S. epidermidis* during skin colonization

Our genetic method provides a powerful new tool to probe the mechanisms by which *S. epidermidis* interacts with host immune cells during colonization. In order to inform a candidate gene approach for immune modulatory microbial ligands, we first set out to identify the host receptors required for an adaptive immune response to *S. epidermidis*, using a previously established system in which *S. epidermidis* is applied topically and immune cells are isolated and phenotyped (**Figure 2A**) (Naik et al., 2012).

**Fig. 2.**
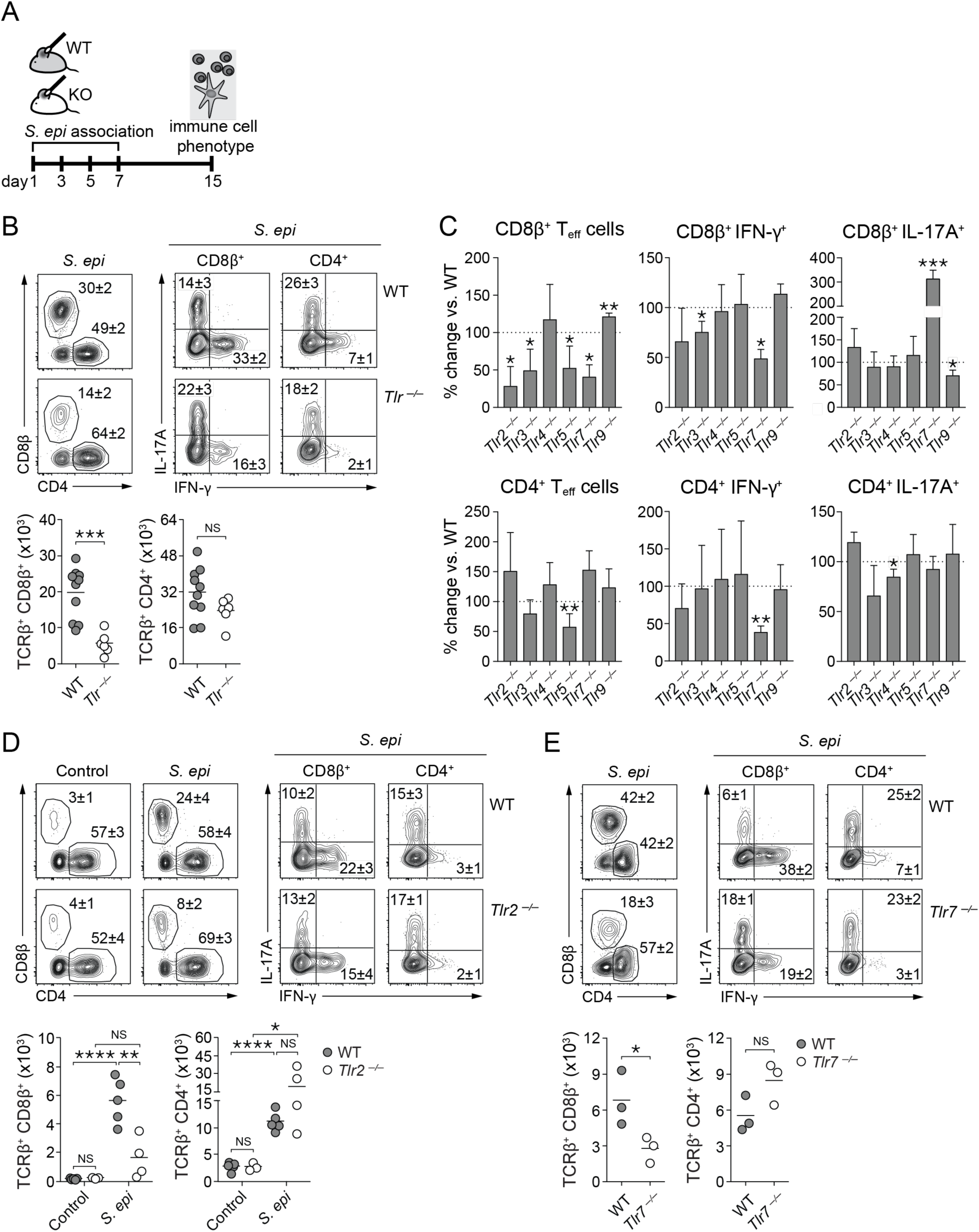
*S. epidermidis* activation of TLR signaling induces distinct T cell populations. (A) Experimental schematic of *S. epidermidis* topical association and collection of ears for skin immune phenotyping. (B) Frequencies (top) of total, IFN-γ^+^ or IL-17A^+^ CD8^+^ or CD4^+^ effector T cells in the skin of *S. epidermidis*-associated wild-type (WT) or complete TLR-deficient (*Tlr^−/−^*). Graphs (bottom) represent the absolute numbers of total CD8^+^ or CD4^+^ effector T cells where each dot represents data from a single mouse. (C) Percentage change in the numbers of skin CD8^+^ or CD4^+^ effector T cells and in the frequencies of IL-17A- or IFN-γ-producing CD8^+^ or CD4^+^ effector T cells in single TLR knockout mice compared to wild-type after *S. epidermidis* association. Each bar represents data from at least 4 mice; data from individual mice are shown in Figure S1D-E. (D-E) Frequencies (top) of total, IFN-γ^+^ or IL-17A^+^ CD8^+^ or CD4^+^ effector T cells in the skin of *S. epidermidis*-associated wild-type (WT), *Tlr2^−/−^* (D), or *Tlr7^−/−^* mice (E). Graphs (bottom) represents the absolute numbers of total CD8^+^ or CD4^+^ effector T cells in wild-type (gray circle), *Tlr2^−/−^* (D, white circle), or *Tlr7^−/−^* mice (E, white circle). “Control” denotes wild-type or *Tlr2^−/−^* unassociated mice. For all figures, NS = not significant, * = p<0.05, ** = p<0.01, *** = p<0.001, and **** = p<0.0001 in an unpaired, two-tailed t-test. In (B), (D) and (E), flow cytometry plots are gated on live CD45.2^+^ CD90.2^+^ TCRβ^+^ Foxp3^−^ cells. Numbers correspond to the frequencies of gated populations ± SD. Data shown are representative of 2-3 independent experiments.

When topically associated onto wild-type specific pathogen free (SPF) mice without any barrier breach or inflammation, *S. epidermidis* strain LM087 promote the accumulation of IFN-γ and IL-17A- producing CD8^+^ (Tc1, Tc17) and CD4^+^ T cells (Th1,Th17), as previously described (**Figure 2B**) (Harrison et al., 2019; Linehan et al., 2018; Naik et al., 2012, 2015). This homeostatic immune response protects against infection by diverse pathogens, including *Leishmania major* and *Candida albicans* (Naik et al., 2012, 2015), and promotes tissue repair after injury (Linehan et al., 2018). However, the *S. epidermidis* ligands and host receptors responsible for triggering this response and differentially promoting distinct classes of T cells remain unknown. Therefore, we set out to test the contribution from different classes of pattern recognition receptors.

In a complete Toll-like receptor deficient mouse (*Tlr-/-*)(Sivick et al., 2014), *S. epidermidis* colonization was impaired in its ability to induce CD8^+^ T cells but was still able to stimulate CD4^+^ T cells (**Figure 2B, S1A**). To determine which specific Toll-like receptors (TLRs) may be involved, we screened a panel of individual mouse TLR knockouts (**Figure 2C-E, S1B-E**). As expected, deletion of *tlr4* and *tlr9* did not decrease the skin T cell response, since their cognate ligands (lipopolysaccharide and viral RNA, respectively) are not present in *S. epidermidis*. In contrast, deletion of *Tlr2*, *Tlr3*, or *Tlr7* resulted in a reduction of the characteristic CD8^+^ T cell response, while the CD4^+^ T cell response remained unchanged (**Figure 2C-E, S1B-E**). On the other hand, deletion of *Tlr5* reduced the CD4^+^ T cell response specifically (**Figure 2C, S1-D**). Interestingly, in the *Tlr3-/-* and *Tlr7-/-* mice, IFN- γ-producing CD8^+^ T cells were dominantly affected, whereas in the *Tlr2-/-* mice, although CD8^+^ T cell responses were decreased, the ability of these cells to produce IFN- γ- or IL-17A-producing was unaffected (**Figure 2D-E, S1C-E**). These results suggest that specific TLRs do not just broadly sense *S. epidermidis* under conditions of colonization; distinct TLRs specifically control CD8^+^ versus CD4^+^ T cell responses and also differentially tune the cytokine polarization and function of T cells in the skin.

We next considered C-type lectin receptors (CLRs), a family of >100 glycan receptors thought to sense diverse fungal and bacterial molecules, including fungal β-glucan (del Fresno et al., 2018), *Mycobacterium tuberculosis* trehalose dimycolate (Ishikawa et al., 2009), and *Helicobacter pylori* lipopolysaccharide (Devi et al., 2015). Given that the *Staphylococcus* cell surface is rich in glycans and glycolipids, we set out to test whether CLRs are involved in the immune response to *S. epidermidis*. We found that deletion of *Clec7a* (Dectin-1) decreased CD8^+^ T cell stimulation without affecting CD4^+^ T cell stimulation, similarly to deletion of *Tlr2* (**Figure 3A-B**). Deletion of *Clec7a* also did not change the balance of IFN-γ and IL-17A secretion by T cells, similarly to deletion of *Tlr2* (**Figure 3B**). In mice lacking both TLR2 and Dectin-1, the CD8^+^ T cell response was significantly decreased but the CD4^+^ T cell response remains unaffected, similarly to the single knockouts (**Figure 3C**). Therefore, either distinct pattern recognition receptors are important for the CD4^+^ T cell response, or that CD4^+^ T cell stimulation relies on more redundant sensing of microbial ligands. In both *Clec7a-/-* and *Tlr2*-/- mice, we also observed a compensatory increase in γδ*-*T cell stimulation by *S. epidermidis* (**Figure S2A-B**). Blocking this γδ-T cell response did not alter the effect of TLR2 or CLEC7A on CD8^+^ T cell stimulation, suggesting that the αβ-T cell response to colonizing *S. epidermidis* is independent from the γδ-T cell response (**Figure S2C-D**). Instead, the γδ-T cell response may be a secondary effect when the αβ-T cell response is unable to be stimulated.

**Fig. 3.**
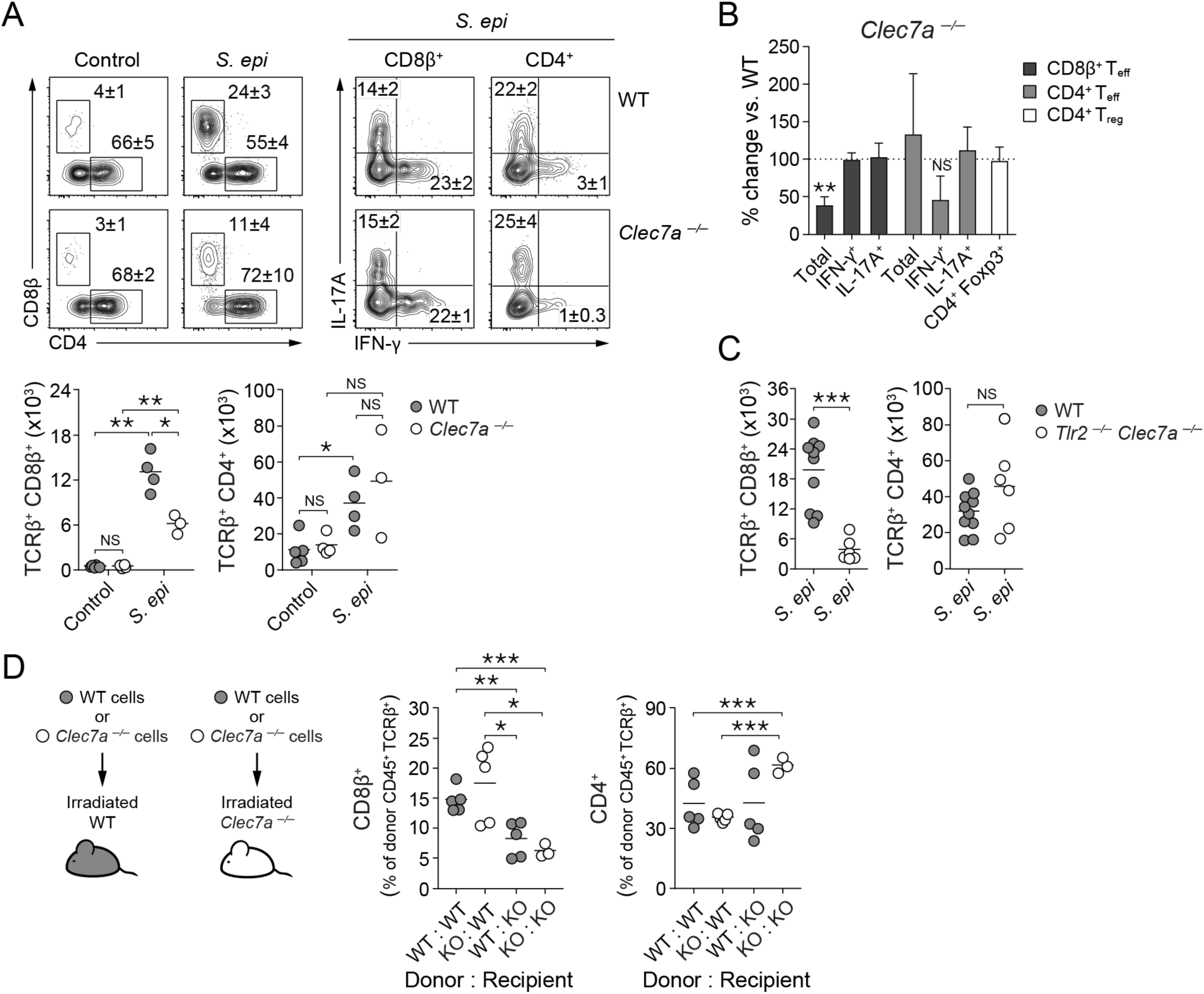
Sensing of *S. epidermidis* by Dectin-1 is required for the CD8^+^ T cell colonization response. (A) Frequencies (top) of total, IFN-γ^+^ or IL-17A^+^ CD8^+^ or CD4^+^ effector T cells in the skin of wild-type or Clec7a^−/−^ mice previously associated with *S. epidermidis* or unassociated (Control). Graphs show the numbers of skin CD8^+^ or CD4^+^ effector T cells in unassociated (Control) or *S. epidermidis*-associated wild-type (WT, grey circles) or *Clec7a*-/- mice (white circles) at day 15. Flow cytometry plots are gated on live CD45.2^+^ CD90.2^+^ TCRβ^+^ Foxp3^−^ cells and the numbers in flow cytometry plots correspond to the frequencies of gated populations ± SD. (B) Percentage change of the number of skin total CD8^+^ or CD4^+^ effector T cells, of the frequencies of IL-17A- or IFN-γ-producing CD8^+^ or CD4^+^ effector T cells, and of the number of regulatory T cells in *Clec7a-/-* mice compared to wild-type after *S. epidermidis* association. Each bar represents data from at least 5 mice. (C) Absolute numbers of skin total CD8^+^ or CD4^+^ effector T cells in *S. epidermidis*-associated wild-type (grey circles) or *Tlr2^−/−^Clec7a*^−/−^ double knockout mice (white circles). (D) Frequencies of skin CD8^+^ or CD4^+^ effector T cells within the donor hematopoietic cell population in *S. epidermidis*-associated bone marrow chimeric mice with either wild-type donor bone marrow (grey circles) or *Clec7a*-/- donor bone marrow (white circles). Nomenclature of mice is donor genotype:recipient genotype. Data showing cytokine production by these T cells is in Figure S2D.

### Sensing of *S. epidermidis* by epithelial and hematopoietic cells within the skin

During bacterial colonization of the skin surface, immune sensing can be mediated by epithelial cells, such as keratinocytes or neurons, or by traditional antigen-presenting cells, such as dendritic cells (Kashem et al., 2015; Lai et al., 2009; Naik et al., 2015). To determine whether the expression of CLEC7A on the epithelial or the hematopoietic compartment in the skin is responsible for driving CD8^+^ T cell stimulation by *S. epidermidis*, we generated bone-marrow chimeras in wild-type or *Clec7a*-/- knockout mice (**Figure 3D**). Surprisingly, replacing the myeloid compartment in wild-type with donor *Clec7a*-/- cells did not alter either the CD8^+^ or CD4^+^ T cell response to *S. epidermidis* colonization (**Figure 3D, S2E**). In contrast, *Clec7a*-/- recipient mice with a bone marrow transplant from wild-type mice phenocopied the loss of CD8^+^ T cell stimulation observed in the *Clec7a*-/- knockout (**Figure 3D, S2E**). Although Langerhans cells are a radiation-resistant population of immune cells in the skin, they likely do not represent an important component to the induction of CD8^+^ T cell stimulation (Naik et al., 2015). These data suggest that although a specific population of CD103^+^ dendritic cells are required for induction of CD8^+^ T cell responses (Naik et al., 2015), C-type lectin receptor sensing of microbial molecules most likely by tissue-resident non-myeloid cells is also required.

### *S. epidermidis* cell envelope mutants induce distinct changes to adaptive immune response

In light of our observation that host TLR2 and CLEC7A signaling are required for the CD8^+^ T cell response to colonizing *S. epidermidis*, we hypothesized that the relevant microbial ligands for the induction of these cells would be lipids, glycolipids, or glycans, which are found in abundance on bacterial cell surfaces (Blanc et al., 2013; Brown et al., 2013). Using the genetic method described above (**Figure 1A**), we created a panel of mutants of the primary human isolate *S. epidermidis* LM087 with defined alterations in cell surface structures. Based on homology to well-described genes in *S. aureus* (**Table S1**), we made deletions in *S. epidermidis* that selectively removed wall teichoic acid (Δ*tagO*), the D-alanine modification on wall and lipoteichoic acid (Δ*dltA*), lipoproteins (Δ*lgt*), and cell-wall associated proteins (Δ*srtA*) (**Figure 4A, S3A-B**). We observed that all mutants were capable of colonizing mouse skin at the time of immune phenotyping (**Figure 4B**). Although we observed, minor variability in the level of colonization between the various mutants, there was no significant correlation between CFU count and CD8^+^ or CD4^+^ T cell responses (**Figure S4D**).

**Fig. 4.**
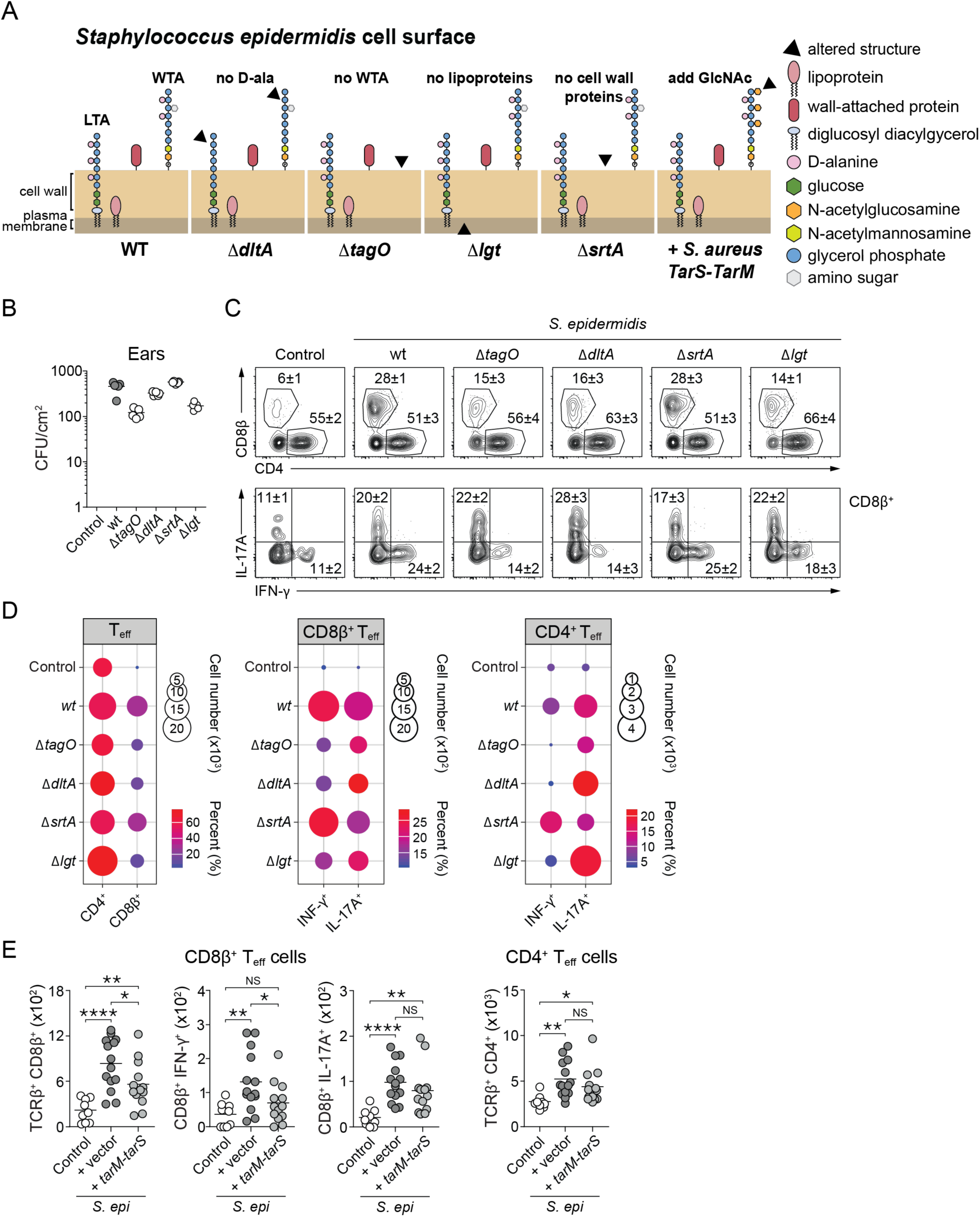
*S. epidermidis* cell envelope mutants yield distinct patterns of immune cell induction. (A) Schematic showing structural alterations in the cell envelope of *S. epidermidis* mutants. WTA = cell-wall attached wall teichoic acids, LTA = plasma-membrane anchored lipoteichoic acids (LTA). (B) Enumeration of *S. epidermidis* colony-forming units (CFU) isolated from mouse skin on day 14 after application of wild-type or mutant *S. epidermidis* strains (*n* = 4-5 per group). (C) Frequencies of skin total CD8^+^ and CD4^+^ effector T cells (top) and IFN-γ- or IL-17A-producing CD8^+^ effector T cells (bottom) in wild-type mice that were unassociated (Control) or associated with wild-type or mutant *S. epidermidis*. Flow plots are gated live CD45.2^+^ CD90.2^+^ TCRβ^+^ Foxp3^−^ cells and the numbers in flow cytometry plots correspond to the frequencies of gated populations ± SD. (D) Mean of absolute numbers (represented by circle size) and frequencies (represented by the circle color) of total, IFN-γ^+^ or IL-17A^+^ CD8^+^or CD4^+^ effector T cells isolated from the skin of mice that were unassociated (Control) or associated with wild-type or mutant *S. epidermidis* strains. Data is aggregated from at least 4 mice per group and representative of multiple experimental replicates. Raw data is in Figure S4. (E) Absolute numbers of total CD4^+^ effector T cells and of total, IFN-γ^+^ or IL-17A^+^ CD8^+^ effector T cells in wild-type mice that were unassociated (Control) or associated with *S. epidermidis* containing empty vector pLI50 (+vector) or *S. epidermidis* expressing *S. aureus* TarM and TarS on pLI50 (+*tarM-tarS*).

Eliminating *S. epidermidis* lipoproteins and D-alanylation of teichoic acids (Δ*lgt* and Δ*dltA*, respectively) almost completely abolished CD8^+^ T cell induction without a significant effect on CD4^+^ T cell stimulation (**Figure 4C-D, S3C-D**), similar to the deletion of *Tlr2* and *Clec7a* in the host. In contrast, the removal of wall teichoic acids (Δ*tagO*) significantly impaired both CD8^+^ and CD4^+^ T cell accumulation (**Figure 4C-D, S3E**). The loss of T cell responses in the Δ*tagO* mutant was intriguing in light of two related observations. First, *S. epidermidis* LM087 is capable of stimulating CD8^+^ T cells, whereas *S. aureus* NCTC8325 is not (Naik et al., 2015). This raises the possibility that a molecular difference in the cell envelopes of these two strains could contribute to their differential capacity to stimulate immune signaling. Second, *S. aureus* and *S. epidermidis* teichoic acids are predicted to vary in a specific way. *S. aureus* harbors multiple glycosyltransferases that are not found in *S. epidermidis*; these include TarM and TarS, which attach *N*-acetylglucosamine (GlcNAc) moieties to the repeating unit of wall teichoic acid (Sobhanifar et al., 2015, 2016; Xia et al., 2010) (**Table S1**).

We hypothesized that some of these GlcNAc moieties may block immune recognition of *S. aureus* wall teichoic acid. To explore this possibility, we engineered *S. epidermidis* to express TarS and TarM on a medium-copy plasmid; mice colonized with this mutant exhibit decreased CD8^+^ and unchanged CD4^+^ T cell stimulation (**Figure 4E**). The effect is less profound than complete deletion of WTA (Δ*tagO*) or D-alanine modification of teichoic acids (Δ*dltA*), perhaps due to incomplete or inefficient glycosylation of *S. epidermidis* teichoic acids by TarS/TarM. But consistent with the deletion of *tagO* and *dltA*, addition of *tarS* and *tarM* disproportionately affected IFNγ-producing CD8^+^ T cells over IL17A-producing CD8^+^ T cells (**Figure 4E**). Together, these data suggest that sensing glycan moieties on the *S. epidermidis* surface modulates CD8^+^ T cell stimulation, with a disproportionate effect on IFN-γ production.

These data also support the idea that *S. aureus* may have evolved to mask its teichoic acids to avoid these commensal T cell responses, thus leading to increased escape from immune surveillance and increased virulence as well as a loss of the beneficial host homeostatic response. A recent report shows that methicillin-resistant *Staphylococcus aureus* strains often contain an alternative wall teichoic acid glycosyltransferase, TarP, encoded in prophages that elicit much lower levels of immunoglobulin G and lead to pathogen evasion of humoral immunity (Gerlach et al., 2018). Here, we show an effect of differential teichoic acid glycosylation on cellular immunity. While pathogens like *S. aureus* have altered their immune epitopes to escape immune detection, commensal microbes like *S. epidermidis* may have differentially evolved the same structures to be able to colonize effectively and provoke a measured and specific immune response that benefits host epithelial health and immune barrier function.

Finally, the loss of cell-wall associated proteins (Δ*srtA*) did not affect the adaptive immune response to *S. epidermidis* under conditions of colonization (**Figure 4C**). Sortase substrates are critical for *S. aureus* pathogenesis (Mazmanian et al., 2000) and infection-associated inflammation in models of infection, such as septic arthritis (Jonsson et al., 2002). These data further highlight that immune responses to *S. aureus* and *S. epidermidis* are triggered by distinct molecular components of the *Staphylococcus* cell wall.

### *S. epidermidis* mutants yield divergent outcomes in colonization versus intradermal injection

In light of our finding that *S. epidermidis* teichoic acids and lipoproteins are critical to immune signaling during colonization, we next asked whether these molecules are required in a different context: the immune response to *S. epidermidis* during skin invasion. To our surprise, when *S. epidermidis* mutants deficient in wall teichoic acids, teichoic acid D-alanylation, and lipoprotein biogenesis were injected intradermally into the ear, we found that all mutants could robustly induce CD8^+^ or CD4^+^ T cell responses to similar levels as wild-type *S. epidermidis* (**Figure 5B, S5**), in contrast to the case of topical association. However, there were significant alterations in the stimulation of innate immune populations (**Figure 5D**). On the other hand, while the absolute number of T cells was unchanged, the quality of type 1 responses by both CD4^+^ and CD8^+^ T cells were significantly decreased with Δ*tagO* and Δ*dltA* but not with Δ*lgt* (**Figure S5**). Further, the loss of D-alanylation of teichoic acids (Δ*dltA*) decreased neutrophil infiltration but stimulated the other innate immune populations (**Figure 5C**). Taken together, these results support the idea that in the context of tissue damage and inflammation, teichoic acids and lipoproteins have a distinct impact on innate inflammatory responses. These data underscore the importance of context (e.g., state of the barrier, tissue site of sensing) to the host immune response to microbes, and they are consistent with a model in which teichoic acids and lipoproteins serve as colonization factors that are either dispensable (for T cell responses) and differentially interpreted (for innate cell inflammatory infiltrate) by the host in the context of pathogenic settings. Thus, our findings suggest that the molecular language of commensalism in the skin is distinct from that of pathogen-immune interactions.

**Fig. 5.**
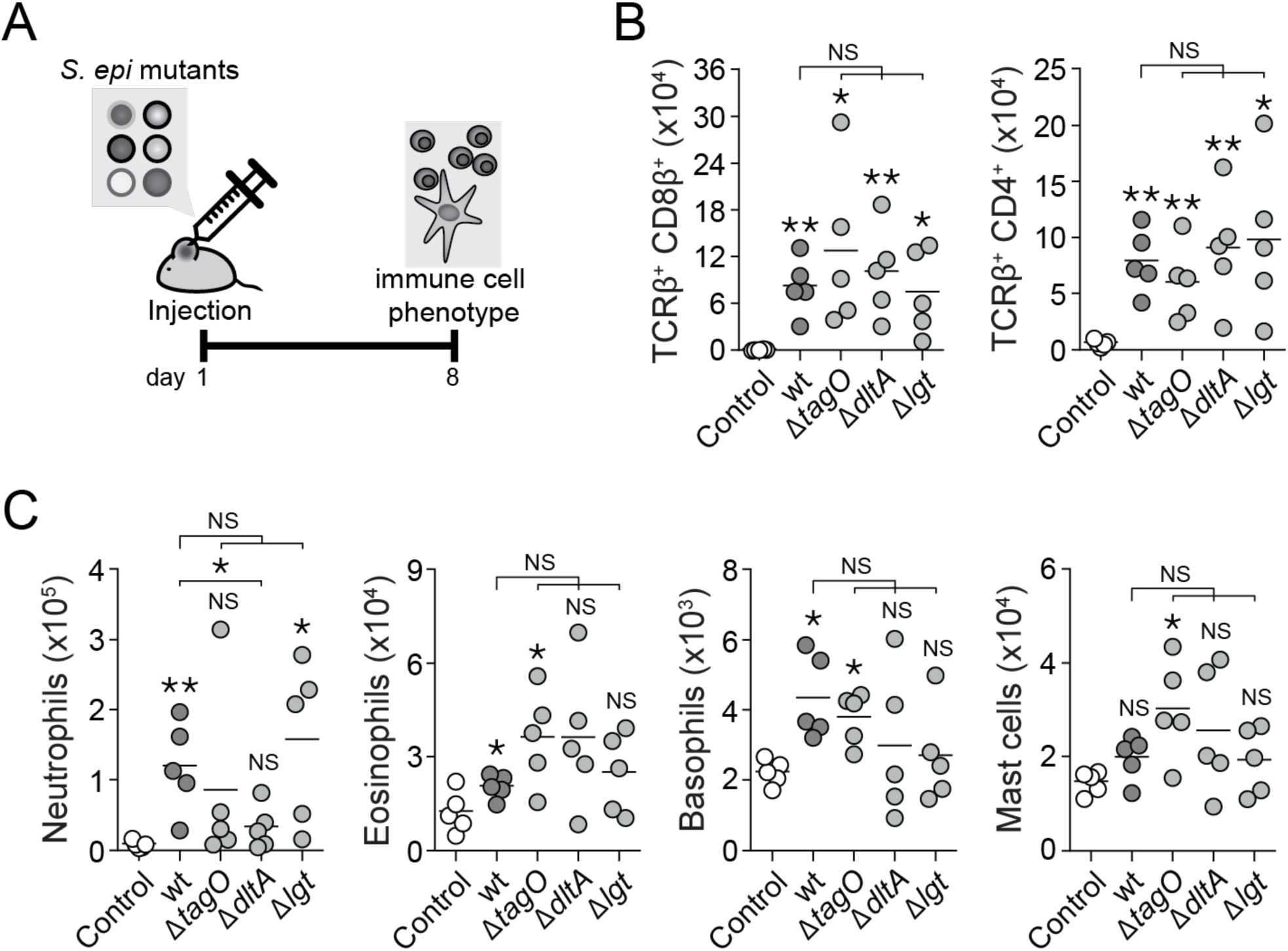
*S. epidermidis* cell envelope mutations alter the innate inflammatory response but not the T cell response in an infection model. (A) Experimental schematic of infection model where *S. epidermidis* is injected intradermally into the ear on day 1 and immune phenotyping of the skin occurs on day 8. (B) Absolute numbers of CD8^+^ and CD4^+^ effector T cells after intradermal injection of wild-type or mutant *S. epidermidis*. (C) Absolute numbers of innate immune populations in unassociated mice (Control, white circles) or 1 week after intradermal injection of wild-type (wt, dark grey circles) or mutant (light grey circles) *S. epidermidis.* Gating strategy for innate cell populations is shown in Figure S4. For all figures, each dot represents a single mouse. NS = not significant, * = p<0.05 and ** = p<0.01 in an unpaired, two-tailed t- test. Asterisks directly above each set of data points denotes comparison to the unassociated mice (Control, white circles). Data shown are representative of 2 independent experiments.

## DISCUSSION

Hundreds of bacterial species colonize mammalian barrier sites. Of these, *S. epidermidis* is among the most prevalent: it is present on every human and is typically the single most abundant bacterial species on skin. Given its predominance in the human microbiome, remarkably little is known about the mechanisms by which *S. epidermidis* interacts with the host, due in large part to the inability to genetically manipulate most strains of *S. epidermidis*. This is a challenge common to the study of nearly all of the most common members of the human microbiota. Previous efforts to study *S. epidermidis*-host interactions have focused on characterizing the immune cells that sense *S. epidermidis* colonization (Lai et al., 2009; Linehan et al., 2018; Naik et al., 2012, 2015). However, the lack of a reliable genetic system for *S. epidermidis* has precluded mechanistic work on the bacterial components of microbiome-host interactions during skin colonization.

The genetic method we introduce here opens the door to discovering the *S. epidermidis* molecules that govern its interactions with the host. Electroporation is a versatile method of introducing foreign DNA into bacteria; the factors that enabled us to apply it to *S. epidermidis* include outgrowth in media containing hyperosmolar sorbitol, harvesting cells in a narrow window during late exponential phase, stringent washing with a large volume of 10% glycerol to remove salts, adding a heat shock before or after electroporation (Dorella et al., 2006; Löfblom et al., 2007), and use of a methylase-deficient *E. coli* strain to produce the donor DNA (Monk et al., 2012). With modest efforts to optimize heat shock time and temperature, these techniques can very likely be generalized to other Gram-positive bacteria that are currently genetically intractable. Notably, these techniques could make it easier to study primary commensal isolates, which have different features than more commonly studied model organisms.

Another strength of our approach is that our experiments involve colonizing native mouse skin with live bacteria, so our immune readout integrates the *in situ* epithelial and immune signals in their native structure. Previous efforts have characterized purified microbial components in cell culture or in assays of *S. epidermidis* invasion, but not in the context of colonization (Deininger et al., 2003; Sabaté Brescó et al., 2017; Volz et al., 2018). Our data reveal the role of individual *S. epidermidis* ligands, presented by a colonist through an intact barrier, in triggering specific components of the adaptive immune response. Notably, teichoic acids and lipoproteins have previously been implicated in *S. aureus* pathogenesis (Brown et al., 2012; Hashimoto et al., 2006; Stoll et al., 2005). Their role in commensal immune sensing indicates that similar molecules can elicit highly distinct host responses depending on context (e.g., the site of sensing and the identity of the presenting microbe). Although *S. epidermidis* teichoic acids and lipoproteins are critical for CD8^+^ and/or CD4^+^ T cell responses in the skin during colonization, these molecules are surprisingly dispensable for the anti-pathogen immune response when *S. epidermidis* is injected intradermally. Instead, different *S. epidermidis* cell surface alterations changed the response of distinct innate immune populations. Intradermal injection bypasses the epithelial barrier and disrupts the structural organization of skin immune system. Thus, not only are the microbial molecules that drive immune responses distinct between commensal and pathogenic host contexts, but their interpretation by the immune system and how innate immune signals are transduced to adaptive immune signals are also distinct. The divergence in immune response to the same set of *S. epidermidis* mutants in different contexts of microbial sensing also underscores the difficulty in drawing direct comparisons between *in vitro* results co-culturing bacteria with immune cells to *in vivo* responses. Importantly, our data show that the complex glycans and lipoglycans in the cell wall of *S. epidermidis* may have evolved to serve as “commensalism factors” rather than virulence factors.

Similarly, we show that TLRs and CLRs, host receptors described predominantly as sensors of microbial “danger signals” in the context of pathogen invasion, are critical for sensing commensals under homeostatic conditions of skin colonization. TLR2 was previously identified as the receptor for *Staphylococcus* lipoproteins and lipoteichoic acids that transduces a pro-inflammatory signal (Blanc et al., 2013; Kurokawa et al., 2009, 2012; Schröder et al., 2003). TLR2 was not previously known to participate in commensal-induced tissue protective responses in the skin. Additionally, lipoproteins and lipoteichoic acids can stimulate immune cells through TLR1-, 2-, and 6-independent mechanisms that are not well-defined (Buwitt-Beckmann et al., 2006; Kaesler et al., 2016). We now show that in addition to TLR2, CLRs including (but not limited to) Dectin-1 are required for T cell stimulation by *S. epidermidis*. Our data complements recent reports that *S. aureus* wall teichoic acids can be sensed by another CLR, Langerin, on Langerhans cells in the skin to induce inflammation (Dalen et al., 2019; van Dalen et al., 2018). Thus, in parallel to the gut, where TLR stimulation by commensal microbiota induces a tissue-protective response (Rakoff-Nahoum et al., 2004), our data demonstrate that TLR and CLR signaling in response to a multiple *S. epidermidis* surface molecules is also required for a tissue-protective adaptive immune response in the skin.

Previous literature has shown that CLR and TLR signaling pathways can interact either positively or negative to generate a combined immune output (del Fresno et al., 2018). For example, the CLR DC-SIGN inhibits TLR4 signaling to block LPS-induced dendritic cell maturation (Doz et al., 2007; Gringhuis et al., 2007). On the other hand, CLEC7A and TLR2 have been shown to cooperate within the same cell to increase inflammatory cytokine production (Gantner et al., 2003). In our case, both the single and double knockouts of host *Tlr2* and *Clec7a*, the CD8^+^ T cell response to colonizing *S. epidermidis* was dampened but not completely abolished, while the CD4^+^ T cell response was largely unaffected. These suggest that TLR2 and CLEC7A function in parallel to sense colonizing *S. epidermidis* but additional receptors are involved to generate the CD4^+^ T cell response.

Where these pattern recognition receptors are located, and which cell populations are involved to ultimately generate the adaptive immune response to colonizing *S. epidermidis* has also been unclear. Previously, it was shown that antigen presentation by CD103^+^ dendritic cell to CD8^+^ T cells via a non-classical MHC I molecule H2-M3 is required for *S. epidermidis* stimulation of CD8^+^ IL17A-producing T cells during colonization (Linehan et al., 2018; Naik et al., 2015). Here, in addition to antigen-presentation by dendritic cells, we show that *S. epidermidis* activation of the CLR CLEC7A may occur through non-myeloid cells in the skin, and this process is also required to generate the CD8^+^ T cell response. Commensal bacteria were previously known to activate TLR signaling in keratinocytes in the setting of skin injury (Lai et al., 2009). Our data now demonstrate that signaling by non-myeloid skin cells likely also occurs during commensal colonization and this signaling may link the tissue context of commensal sensing to the resultant adaptive immune response.

In sum, we developed new methods that enable us to create a panel of systematic cell surface deletions or alterations in the background of a primary human isolate of *S. epidermidis*. Using a model of commensal skin colonization, we dissect the microbial and host components responsible for the *S. epidermidis-*immune communication. Specifically, we show that not only TLRs, but also CLRs and non-myeloid skin cells, are critical to triggering the homeostatic T cell response to *S. epidermidis* colonization. Our data demonstrate that commensalism and pathogenesis are sensed by a common molecular language, but the host context in which the signal is received governs the inflammatory outcome. Taken together, our work opens the door to a broader and more mechanistic exploration of *S. epidermidis*-host interactions with a wide variety of primary human *S. epidermidis* isolates. This more mechanistic understanding can in turn provide novel therapeutic targets and methods for modulating immune activation in inflammatory skin diseases.

## METHODS

### Mice

Wild-type (WT) C57BL/6 specific pathogen-free mice were purchased from Taconic Farms. B6.[KO]TLR3 N10 (*Tlr3* ^−/−^), C57BL/6J-CD45a (CD45.1 WT), C57BL/6J × B6.SJL-CD45a/Nai F1 (CD45.1/CD45.2 WT) mice were obtained through the NIAID-Taconic exchange program. B6.129S6- *Clec7a^tm1Gdb^*/J (*Clec7a* ^−/−^) mice and their C57BL/6J wild-type controls were purchased from The Jackson Laboratory. B6.129S1-*Tlr5^tm1/Flv^*/J (*Tlr5* ^−/−^) and B6.129P2-*Tlr9^tmAki^* (*Tlr9* ^−/−^) mice were kindly provided by Dr. G. Trinchieri (National Cancer Institute/NIH). B6.129P2-*Tlr2^tmAki^* (*Tlr2* ^−/−^) and B6.129P2-*Tlr4^tm1Aki^* (*Tlr4* ^−/−^) mice backcrossed 11 generations onto the C57BL/6 (Taconic) background were generous gifts from Dr. Alan Sher (NIAID/NIH). B6.129S1-*Tlr7^tm1Flv^*/J (*Tlr7* ^−/−^) and B6.129S6-*Card9^tm1Xlin^*/J (*Card9* ^−/−^) mice were generous gifts from Dr. S. Bolland (NIAID/NIH) and Dr. M. Lionakis (NIAID/NIH), respectively. *Tlr2* ^−/−^*Clec7a* ^−/−^ and *Tlr2* ^−/−^*Card9* ^−/−^ mice were generated by crossing the F1 progeny of *Tlr2* ^−/−^ ×*Clec7a* ^−/−^ breeders and *Tlr2* ^−/−^×*Card9* ^−/−^, respectively. *Tlr2* ^−/−^*Tlr4* ^−/−^*Tlr5* ^−/−^*Unc93b1^3d/3d^* (*Tlr* ^−/−^) mice were generated by and obtained from Dr. G. Barton (University of California, Berkeley). For experiments with knockout mice, littermate controls were used as wild-type controls, and/or wild-type (littermate or C57BL/6) and knockout mice were co-housed at 3-5 weeks of age for 2-3 weeks in the same cage, then separated prior to the start of experimental manipulations. All mice were bred and maintained under pathogen-free conditions at an American Association for the Accreditation of Laboratory Animal Care (AAALAC)- accredited animal facility at the NIAID and housed in accordance with the procedures outlined in the Guide for the Care and Use of Laboratory Animals. All experiments were performed at the NIAID under an animal study proposal (LISB20E) approved by the NIAID Animal Care and Use Committee or at Stanford University under the animal study protocol (32872) approved by the Stanford Laboratory Animal Care Committee. Sex and age-matched mice between 6 and 12 weeks of age were used for each experiment. When possible, preliminary experiments were performed to determine requirements for samples size, taking in account resources available and ethical, reductionist animal use. In general, each mouse of the different experimental groups is reported. Exclusion criteria such as inadequate staining or low cell yield due to technical problems were pre-determined. Animals were assigned randomly to experimental groups.

### Genetic engineering of Staphylococcus epidermidis

To generate electrocompetent cells, *S. epidermidis* was grown in Difco Brain Heart Infusion media (BD 241810) with 0.5 M sorbitol (BHIS) overnight at 37^°^C with shaking and then back-diluted to an OD of 0.15 to 0.25. Back-diluted cultures were grown in BHIS until an OD of 0.7 to 0.9 at 37^°^C with shaking. At this time, cultures were placed on ice and then pelleted at 3500xg for 10 minutes at 4^°^C. Cultures were resuspended in equal volume 10% ice-cold glycerol. Spin and resuspension steps were repeated for a total of four to five 10% glycerol washes. Each 50 ml of culture OD 0.7 to 0.9 was resuspended in 100 ul of 10% ice cold glycerol and used for one electroporation reaction. Fresh competent cells were made for each electroporation.

Approximately 1 μg of plasmid isolated from DC10B *E. coli* or 1 μg of minicircle plasmid from JCM2 *coli* was added to 100 μl of competent *S. epidermidis* and cells (Johnston et al., 2018; Monk et al., 2012). Method A works for strains LM018, LM061, LM087, LM088, LM095, NIH05001, ATCC12228, SK135, W23144, KPL1815. Method B works for strains ATCC35984 (RP62A), DSM20042, BCM0060. For method A, cells and plasmid in 10% glycerol were heat-shocked at 56^°^C for 2 minutes, then immediately transferred to a 0.1 cm cuvette, electroporated, and transferred to 3 ml of room temperature BHIS. For method B, cells and plasmid in 10% glycerol were pre-warmed at room temperature for 5 minutes, electroporated in a 0.1 cm cuvette, diluted into 1 ml of BHIS prewarmed to 56^°^C to heat-shock for 2 minutes, and then diluted with 3 ml of room temperature BHIS. For both methods, the electroporation program was 2.5 kV, 1 pulse, with a typical time constant of 2.3-2.5 msec using a Bio-Rad Micropulser.

Following heat shock and electroporation, cells in BHIS were recovered at 37°C for 1.5-2 hours if using a replicative plasmid (pLI50, a gift from Chia Lee, Addgene plasmid #13573) and at 28°C for 4 hours if using a temperature-sensitive plasmid (pIMAY, gift from Tim Foster, Addgene plasmid # 68939) (Lee et al., 1991; Monk et al., 2012). After recovery, cells were spun down at 3500×g for 10 minutes and plated onto BHIS plates with chloramphenicol (10 μg/ml). For allelic replacement, transformation plates were grown at 28°C for 2-3 days. At this time, single colonies were grown in BHI/chloramphenicol at 37°C overnight, and serial dilutions were plated onto BHI/chloramphenicol at 37°C overnight to cure the plasmid. Single colonies were then restruck onto BHI/chloramphenicol and plates were grown overnight at 37°C. Colony PCR was done to determine the absence of plasmids. Colonies that successfully lost the plasmid (now integrated into the genome) were grown in BHI without antibiotics at 28 deg for 6-12 hours, and then plated onto BHI/anhydrotetracycline (1 μg/ml) for 2 days to select for double crossover events. Colonies were patched onto BHI/anhydrotetracycline and BHI/chloramphenicol, grown at 37 degrees overnight, and then resultant anhydrotetracycline-resistant but chloramphenicol-sensitive colonies were screened for the appropriate knockout or allelic replacement by colony PCR using flanking primers.

### Topical association and intradermal infection of mice with *Staphylococcus epidermidis*

*Staphylococcus epidermidis* strains (NIHLM087 and mutants made in this background) were cultured for 18 hours in tryptic soy broth at 37°C, not shaking. For topical association of bacteria, each mouse was associated by placing 5 ml of the bacterial suspension, approximately 10^9^ CFU/ml, across the ear skin surface using a sterile cotton swab. Application of bacterial suspension was repeated every other day for a total of four times. In experiments involving topical application of various bacterial species or strains, 18-hour cultures were normalized using OD_600_ to achieve similar bacterial density (approximately 10^9^ colony-forming units/ml (CFU/ml). For some experiments, mice were injected intradermally on day 1 in the ear pinnae with 10^7^ CFU of S. epidermidis strain NIHLM087 and mutants made in this background. For intradermal injections, immune cell phenotype was assayed at day 5.

### Bacteria quantitation

The ear pinnae of topically associated or unassociated control mice were swabbed with a sterile cotton swab previously soaked in tryptic soy broth. Swabs were streaked on Columbia blood agar plates. Plates were then placed at 37°C under aerobic conditions for 18 hours. Colony-forming units (CFU) on each plate were enumerated and the number of CFU was reported per cm^2^ of skin.

### Tissue processing and immune phenotypic analysis

Cells from the ear pinnae were isolated as previously described (Naik et al., 2012). For detection of basal cytokine potential, single cell suspensions were cultured directly *ex vivo* in a 96-well U-bottom plate in complete medium (RPMI 1640 supplemented with 10% fetal bovine serum (FBS), 2 mM L-glutamine, 1 mM sodium pyruvate and nonessential amino acids, 20 mM HEPES, 100 U/ml penicillin, 100 μg/ml streptomycin, 50 mM β-mercaptoethanol) and stimulated with 50 ng/ml phorbol myristate acetate (PMA) (Sigma-Aldrich) and 5 μg/ml ionomycin (Sigma-Aldrich) in the presence of brefeldin A (GlogiPlug, BD Biosciences) for 2.5 hours at 37°C in 5% CO_2_. After stimulation, cells were assessed for intracellular cytokine production as described below. Single cell suspensions were incubated with combinations of the following fluorophore-conjugated antibodies against surface markers:, CD4 (RM4-5), CD8β (eBioH35- 17.2), CD11b (M1/70), CD19 (6D5), CD45 (30-F11), CD45.1 (A20), CD45.2 (104), CD49b (DX5), CD64 (FA-11), CD90.2 (53-2.1), CD117 (2B8), FcεRI (MAR-1), Ly6C (HK1.4), Ly6G (1A8), MHC-II (M5/114.15.2), NK1.1 (PK136), Siglec-F (E50-2440), TCRβ (H57-597), γδTCR (eBioGL3), and TCR Vγ2 (UC3-10A6) in Hank’s buffered salt solution (HBSS) for 20 min at 4°C and then washed. LIVE/DEAD Fixable Blue Dead Cell Stain Kit (Invitrogen Life Technologies) was used to exclude dead cells. Cells were then fixed for 30 min at 4°C using the fixation/permeabilization buffer supplied with the BD Cytofix/Cytoperm (Becton Dickinson) and washed twice with permeabilization buffer. For intracellular staining, cells were then stained with fluorochrome-conjugated antibody against IFN-γ (XMG-1.2), IL-17A (eBio17B7) and Foxp3 (FJK-16s) in permeabilization buffer supplied with the BD Cytofix/Cytoperm kit (BD Biosciences) for 1 hour at 4°C. Each staining was performed in the presence of purified anti-mouse CD16/32 (clone 93) and 0.2 mg/ml purified rat gamma globulin (Jackson Immunoresearch). All antibodies were purchased from eBioscience (Life Technologies), Biolegend, or BD Biosciences. Cell acquisition was performed on an LSR Fortessa or LSR II flow cytometer using FACSDiVa software (BD Biosciences) and data were analyzed using FlowJo software (TreeStar).

### Bone marrow chimera

Recipient mice (*Tlr2* ^−/−^, *Clec7a* ^−/−^ or congenic CD45.1 WT) were given lethal total body irradiation (1,200 rads) and reconstituted the same day intravenously with 10 million bone marrow cells isolated from tibial and femoral bones of *Tlr2* ^−/−^, *Clec7a* ^−/−^ or congenic CD45.1/CD45.2 WT mice. Irradiated and reconstituted mice were given Bactrim (sulfamethoxazole [150 mg/ml] and *N*-trimethoprim [30 mg/ml]) in their drinking water for 2 weeks and switched thereafter to sterile drinking water. Mice were allowed to reconstitute for 6-8 weeks before topical association with *S. epidermidis*.

### γδ-T cell depletion

For γδ T-cell depletion, mice were treated intraperitoneally with 0.5 mg of anti-γδ antibody (clone UC7-13D5), or rat IgG2b isotype control antibody (clone LTF-2, BioXCell) every other day, starting the day before the first association.

### Statistics

Data are presented as mean ± standard error of the mean (SEM) or mean ± standard deviation (SD). Group sizes were determined based on the results of preliminary experiments. Mice were assigned at random groups. Mouse studies were not performed in a blinded fashion. Generally, each mouse of the different experimental groups is reported. Statistical significance was determined using two-tailed unpaired Student’s *t*-test under the untested assumption of normality. All statistical analysis was calculated using Prism software (GraphPad). Differences were considered to be statistically significant when *p*<0.05.

## SUPPLEMENTARY INFORMATION

Figures S1-S5 are available in the online version of the paper.

## ACKNOWLEDGMENTS

We are deeply indebted to members of the Fischbach Group and Belkaid Group for helpful suggestions and comments on the manuscript. We thank Greg Barton (Berkeley) for the generous gift of *Tlr* ^−/−^ mice (unpublished). We thank the National Institute of Allergy and Infectious Diseases (NIAID) animal facility staff. This work was supported by an A.P. Giannini Foundation Postdoctoral Research Fellowship (Y.E.C.); an HHMI Hanna H. Gray Fellowship (Y.E.C.); NIAID Division of Intramural Research (ZIA-AI001115, ZIA- AI001132) (Y.B.); an HHMI-Simons Faculty Scholar Award (M.A.F.); a Fellowship for Science and Engineering from the David and Lucile Packard Foundation (M.A.F.); an Investigators in the Pathogenesis of Infectious Disease award from the Burroughs Wellcome Foundation (M.A.F.); an award from BASF (M.A.F.); an award from the Leducq Foundation (M.A.F.); NIH grants DK110174 (M.A.F.), DK113598 (M.A.F.); the Chan Zuckerberg Biohub (M.A.F.). C.H. is funded by fondation La Roche-Posay, Collège des Enseignants en Dermatologie de France (CEDEF), fondation groupe pasteur mutualité, Société Française de Dermatologie, Philippe Foundation, Fondation pour la Recherche Médicale. This work was supported by the Division of Intramural Research of the National Institute of Allergy and Infectious Diseases (NIAID).

## AUTHOR CONTRIBUTIONS

Y.E.C., N.B., C.H., Y.B., and M.A.F. designed the study and wrote the manuscript. Y.E.C., N.B., C.H., and A.M.M. performed the experiments.

## AUTHOR INFORMATION

M.A.F. is a co-founder and director of Federation Bio. Correspondence and requests for materials should be addressed to and will be fulfilled by the Lead Contact, Michael Fischbach.

**Fig. S1.**
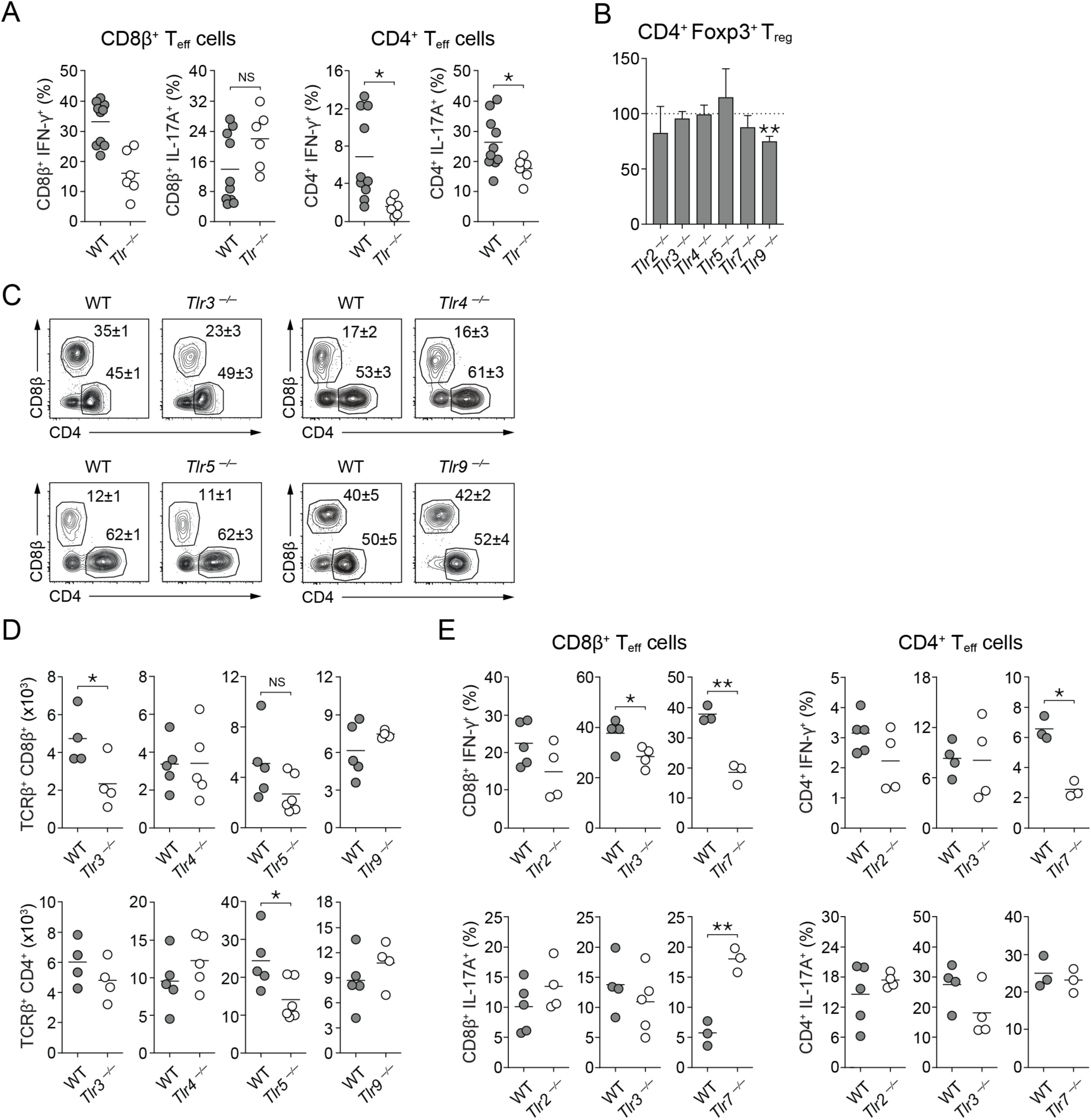
Topical association of *S. epidermidis* in TLR knockout mice. (A) Frequencies of skin IL-17A- or IFN-γ-producing CD8^+^ or CD4^+^ effector T cells in wild-type (WT) or complete TLR knockout (*Tlr*^−/−^) mice that were topically associated with *S. epidermidis*. This is additional data from the same experiment shown in Figure 1B. (B) Percentage change of the frequencies of skin regulatory T cells (TCRβ^+^ CD4^+^ Foxp3^+^) in various single TLR knockout mice compared to wild-type after *S. epidermidis* association. Each bar represents data from at least 4 mice. For panels (C)-(E): Raw data showing (C) frequencies of skin total CD8^+^ and CD4^+^ effectorT cells, (D) absolute numbers of skin total CD8^+^ and CD4^+^ effector T cells, and (E) frequencies of IFN-γ- or IL-17A-producing CD8^+^ or CD4^+^ effector T cells in various single TLR knockout mice after *S. epidermidis* association. In panel C, flow cytometry plots are gated on live CD45.2^+^ CD90.2^+^ TCRβ^+^ Foxp3^−^ cells and the numbers in flow cytometry plots correspond to the frequencies of gated populations ± SD. Panels (B-E) represents additional data from the same experiment shown in Figure 1C-E. For all figures, each dot represents a single mouse. NS = not significant, * = p<0.05 and ** = p<0.01 in an unpaired, two-tailed t-test..

**Fig. S2.**
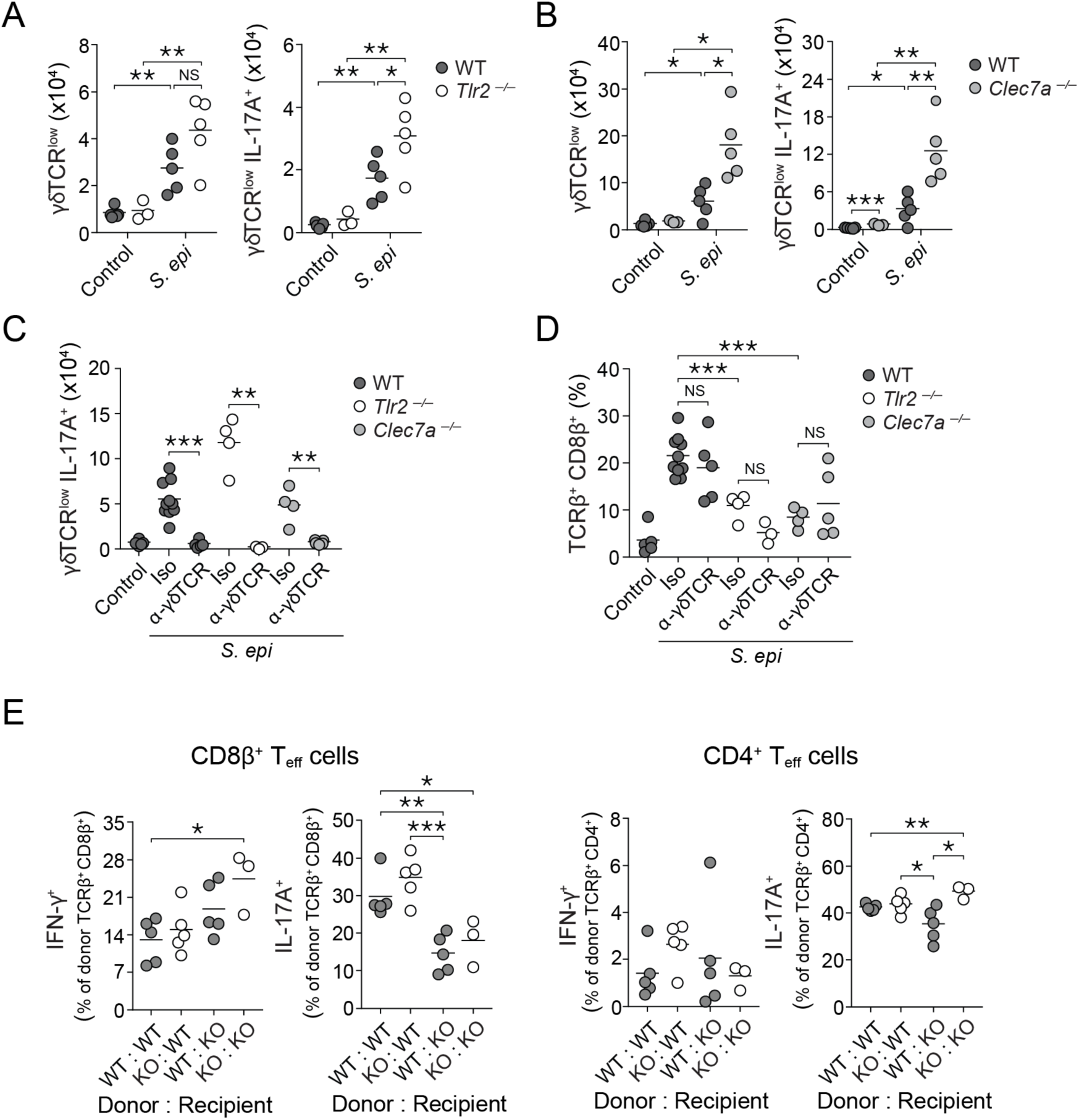
The skin CD8^+^ and CD4^+^ T cell response to *S. epidermidis* association is independent of γδ- T cells. (A-B) Absolute numbers of live CD45^+^ CD90.2^+^ γδTCR^low^ T cells in unassociated (Control) or *S. epidermidis*-associated (*S. epi*) wild-type mice (WT, grey circles), *Tlr2^−/−^* mice (white circles), or *Clec7a^−/−^* mice (light grey circles). (C-D) Wild-type (WT), *Tlr2*^−/−^, or *Clec7a*^−/−^ mice were associated with *S. epidermidis* and treated with an isotype control antibody or an anti-γδTCR antibody. Control animals were untreated and unassociated. (C) Absolute numbers of live CD45^+^ CD90.2^+^ γδTCR^low^ T cells producing IL-17A and (D) frequencies of total CD8^+^ effector T cells in the skin of wild-type (dark circles), *Tlr2*^−/−^ (white circles) or *Clec7a*^−/−^ mice (light grey circles). Panels C-D show data from the same experiment. (E) Frequencies of skin IFN-γ-producing or IL-17A-producing CD8^+^ or CD4^+^ effector T cells within the donor hematopoietic cell population in *S. epidermidis*-associated bone marrow chimeric mice with either wild-type donor bone marrow (grey circles) or *clec7a*-/- donor bone marrow (white circles). Nomenclature of mice is donor genotype:recipient genotype. These are additional data from the same experiment shown in Figure 3D. NS = not significant, * = p<0.05, ** = p<0.01 and *** = p<0.001 in an unpaired, two-tailed t-test.

**Fig. S3.**
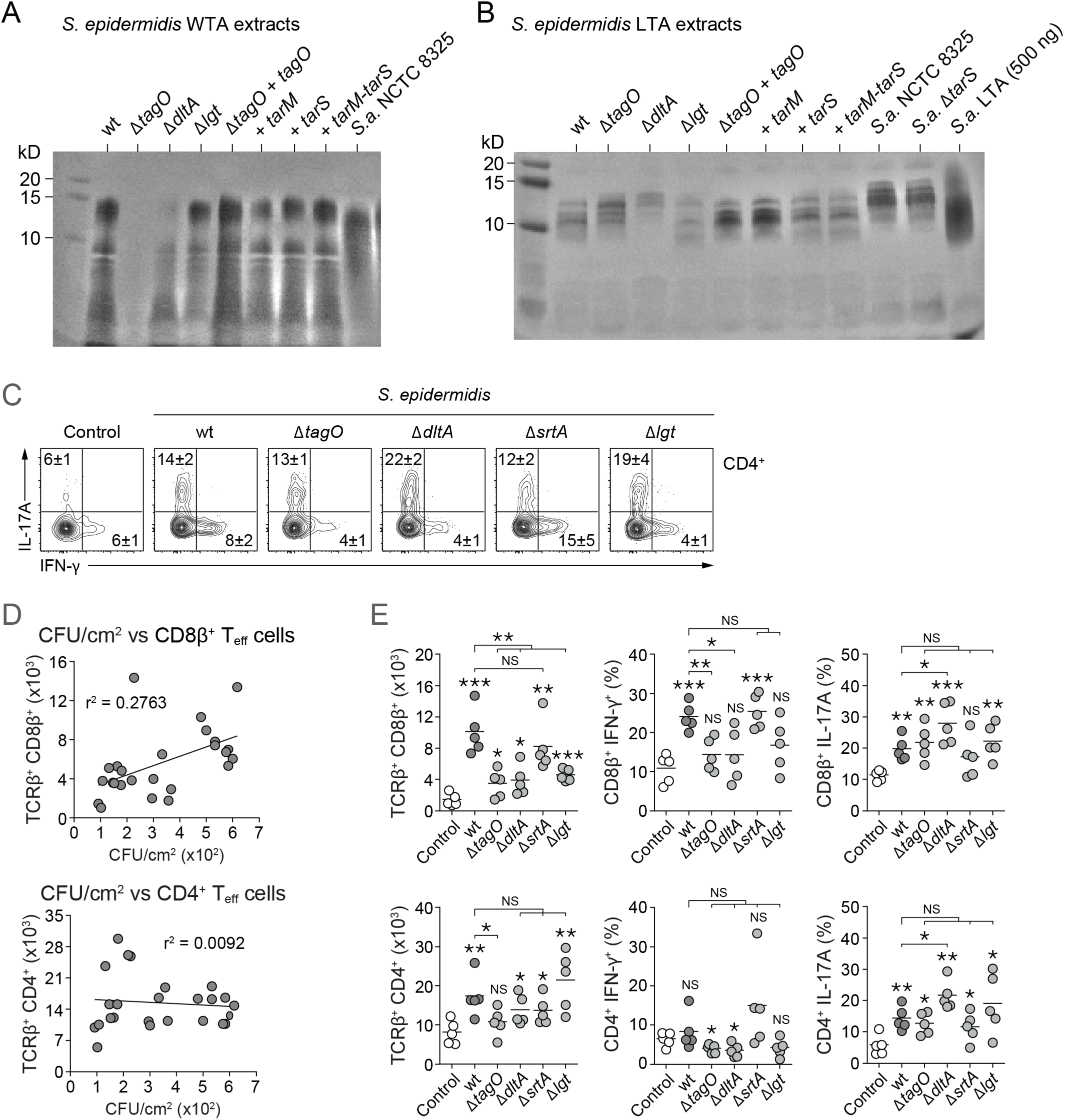
Characterization of *S. epidermidis* mutants. (A) Alcian blue-silver stain of wall teichoic acid extracts from *S. epidermidis* mutants or *S. aureus* strain NCTC8325 using the acid hydrolysis method (Shiratsuchi et al., 2010). As expected *ΔtagO* does not produce wall teichoic acid. (B) Alcian blue-silver stain of lipoteichoic acid, extracted as previously described (Gründling and Schneewind, 2007). The last lane shows 500 ng of commercially available *S. aureus* lipoteichoic acid (Sigma). (C) Frequencies of skin IL-17A^+^ or IFN-γ^+^ CD4^+^ T cells in wild-type mice that were unassociated (Control) or associated with live wild-type or mutant *S. epidermidis*. Flow plots are gated live CD45.2^+^ CD90.2^+^ TCRβ^+^ CD4^+^ Foxp3^−^ cells and the numbers in flow cytometry plots correspond to the frequencies of gated populations ± SD. These are additional data from the same experiment shown in Figure 4C. (D) *S. epidermidis* colonization does not correlate significantly to CD8^+^ or CD4^+^ T cell stimulation. Each dot represents data from one mouse, showing the T cell response on the Y-axis and the number of *S. epidermidis* CFU/cm^2^ recovered by plating from mouse skin swabs at the time of immune phenotyping on the X-axis. (E) Absolute numbers of total CD8^+^ and CD4^+^ effector T cells and frequencies of IFN-γ-, or IL-17A-producing CD8^+^ or CD4^+^ effector T cells in wild-type mice that were unassociated (Control) or associated with live wild-type or mutant *S. epidermidis*. Each dot represents data from one mouse. NS = not significant, * = p<0.05, ** = p<0.01 and *** = p<0.001 in an unpaired, two-tailed t-test. Asterisks directly above each set of data points denotes comparison to the unassociated mice (Control, white circles). Data shown are representative of 3 independent experiments. These are the data used to generate Figure 4D.

**Fig. S4.**
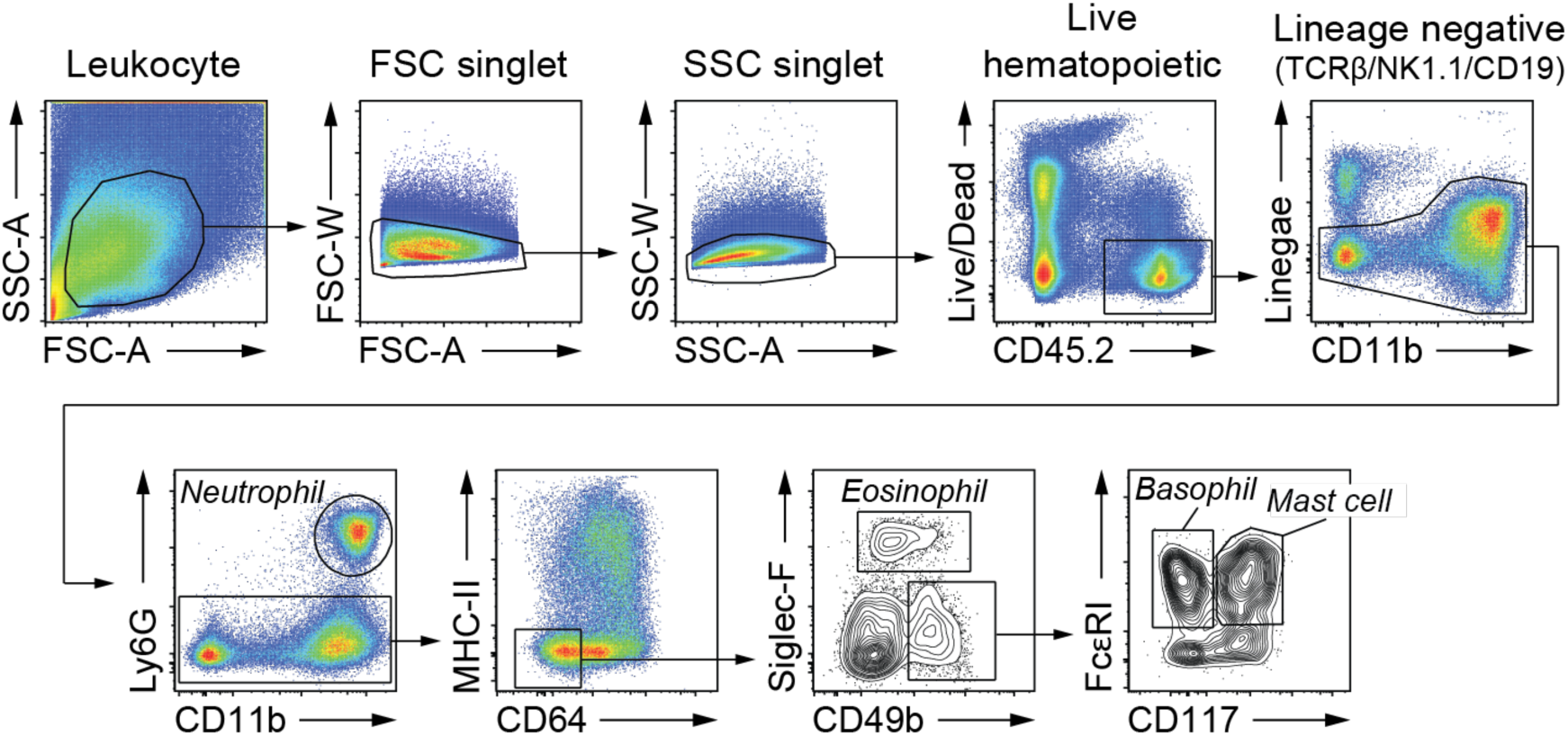
Gating strategy for skin innate immune cells. This gating strategy was used to generate the data shown in Figure 5C.

**Fig. S5.**
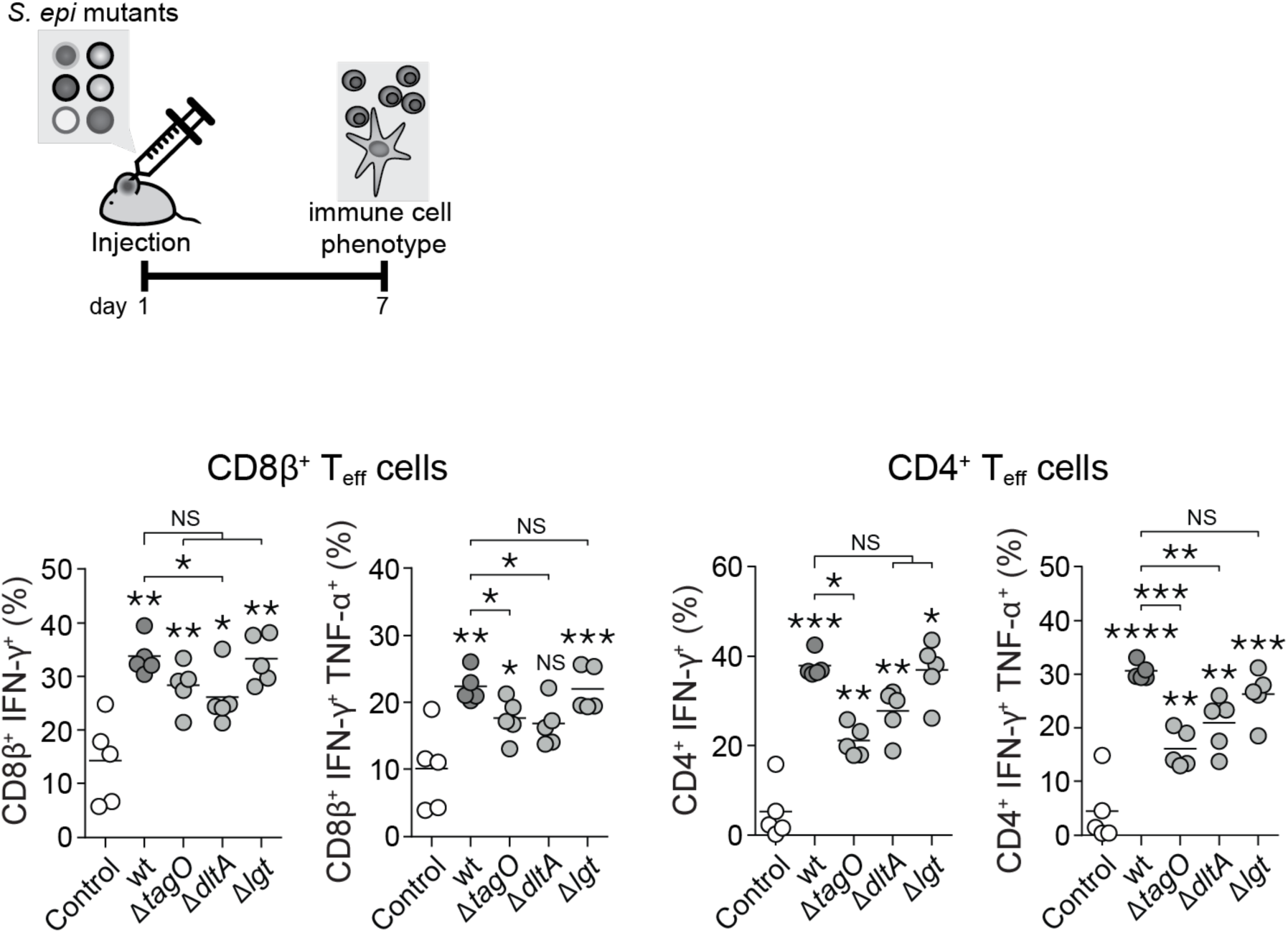
Cytokine production by T cells in response to intradermal injection of wild-type or mutant *S. epidermidis*. Frequencies of IFN-γ^+^ or IFN-γ^+^ TNF-α^+^ CD8^+^ and CD4^+^ effector T cells 1 week after intradermal injection of wild-type or mutant *S. epidermidis*. Each dot represents data from one mouse. NS = not significant, * = p<0.05, ** = p<0.01, *** = p<0.001 and *** = p<0.0001 in an unpaired, two-tailed t-test. Asterisks directly above each set of data points denotes comparison to the unassociated mice (Control, white circles). These are additional data from the same experiment as Figure 5.

**Table S1.**
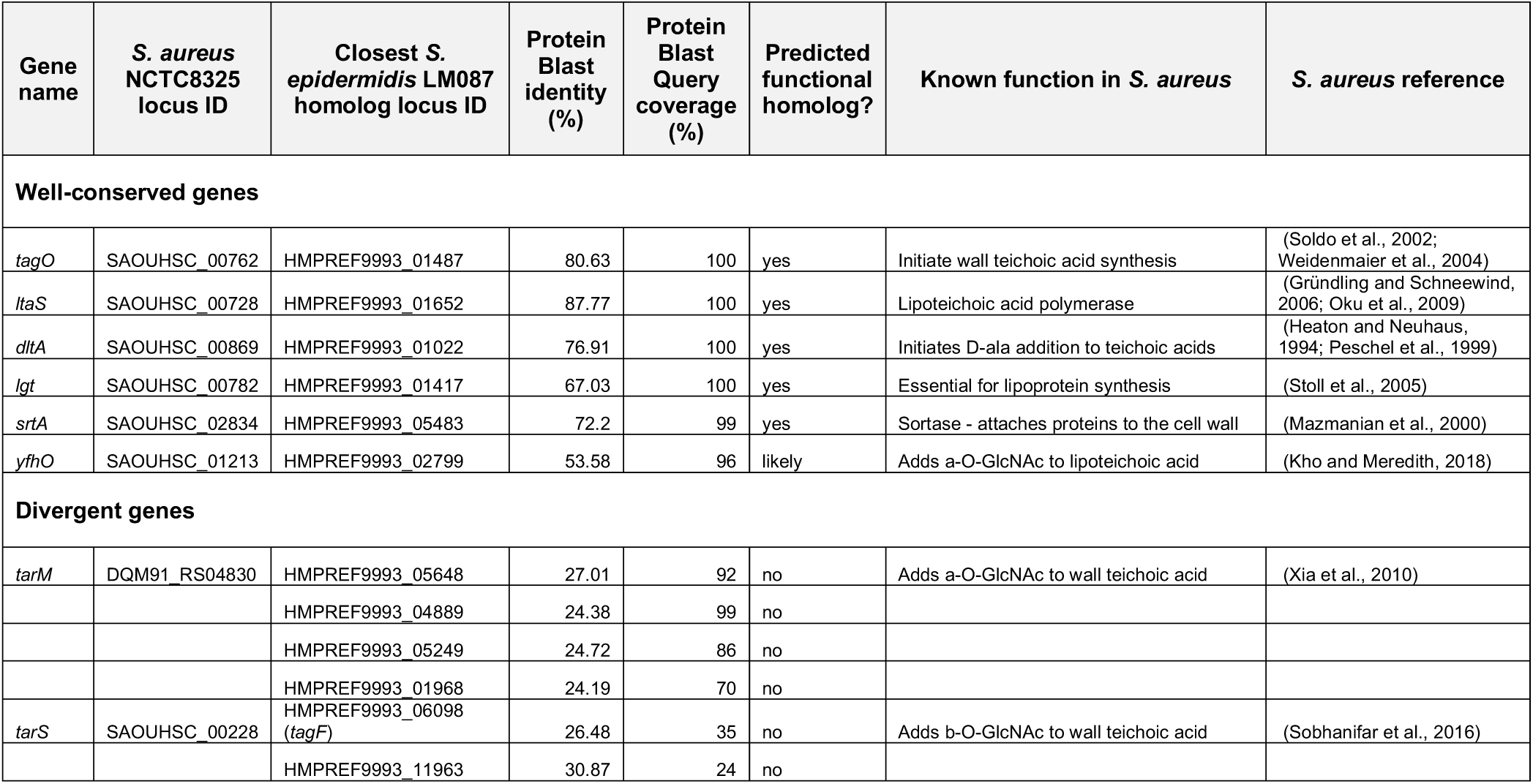

